# PPARγ-dependent and -independent regulation of methionine metabolism by diet-induced obesity and fasting in male mice

**DOI:** 10.64898/2026.03.24.714010

**Authors:** Izabela Hawro, Samuel M. Lee, Rhonda D. Kineman, Jose Cordoba-Chacon

## Abstract

Metabolic dysfunction-associated steatohepatitis (MASH) is associated with increased expression of peroxisome proliferator-activated receptor gamma (PPARγ, *Pparg*) and reduced expression of genes involved in methionine metabolism in the liver. The nuclear receptor PPARγ is activated by fatty acids, and the knockout of *Pparg* in hepatocytes (*Pparg*^ΔHep^) reduced the negative effects of MASH on methionine metabolism. Here, we sought to determine whether hepatocyte *Pparg* is required for the transcriptional regulation of genes involved in hepatic methionine metabolism in conditions with altered fatty acid flux to the liver: fasting, refeeding, and high-fat diet (HFD)-induced obesity/steatosis. Fasting induced liver steatosis and increased the expression of key genes involved in the methionine metabolism in the liver, while 6h-refeeding reversed these effects and reduced the expression of phosphatidylethanolamine N-methyltransferase (*Pemt)* and cystathionine beta synthase (*Cbs)*. Overall, fasting and refeeding did not alter hepatocyte *Pparg* expression nor *Pparg*^ΔHep^ affected fasting and refeeding-mediated regulation of methionine metabolism gene expression. Diet-induced steatosis reduced hepatic *Pemt* expression in control (*Pparg*-intact) mice, and the thiazolidinedione (TZD)-mediated activation of PPARγ in diet-induced obese control (*Pparg*-intact) mice reduced the expression of betaine homocysteine S-methyltransferase (*Bhmt)* and *Cbs*. However, diet-induced steatosis increased hepatocyte *Pparg* expression, and *Pparg*^ΔHep^ blocked the negative effects of HFD and TZD on hepatic methionine metabolism. The PPARγ-dependent reduction of hepatic *Bhmt* and *Cbs* expression was confirmed in mouse primary hepatocytes. Taken together, hepatocyte *Pparg* may serve as a negative regulator of hepatic methionine metabolism in diet-induced obese mice and these actions could contribute to promoting the onset of MASH.

## Introduction

Methionine is an essential sulfur-containing amino acid that is used for protein synthesis or by the methionine adenosyltransferases 1a and 2a (*Mat1a* and *Mat2a*) to generate S-adenosylmethionine (SAM). The SAM-dependent methyltransferases methylate a wide range of substrates including: DNA, lipids and amino acids, to maintain cellular health. For instance, glycine N-methyltransferase (*Gnmt*) uses SAM to convert glycine into sarcosine, and phosphatidylethanolamine N-methyltransferase (*Pemt*) needs three molecules of SAM to synthesize phosphatidylcholine from phosphatidylethanolamine. The transmethylation reactions of SAM produce S-adenosylhomocysteine (SAH), which is then converted into homocysteine by the adenosylhomocysteinease (*Ahcy*). Finally, homocysteine is processed by the cystathionine β-synthase (*Cbs*) in the transsulfuration pathway to produce glutathione, or remethylated by the 5-methyltetrahydrofolate-homocysteine methyltransferase (*Mtr*) or the betaine homocysteine methyltransferase (*Bhmt*) to regenerate methionine (1,2). Of note, the expression of *Mat1a, Gnmt, Pemt, Ahcy, Bhmt*, and *Cbs* is high and almost exclusive to hepatocytes, and most of the whole-body methionine adenosyltransferase activity is found in the liver (1,3-5). In fact, up to 85% of SAM-dependent methylation reactions and nearly 50% of methionine metabolism take place in the liver (6).

Diet-induced obesity and the progression of metabolic dysfunction-associated steatotic liver disease (MASLD) reduce the expression of *Mat1a, Gnmt, Pemt*, nicotinamide N-methyltransferase (*Nnmt*), *Ahcy*, and *Bhmt* in the liver (7-9). In fact, the knockout of *Gnmt, Pemt*, or *Bhmt* leads to the development of steatosis and progression of MASLD to metabolic dysfunction associated steatohepatitis (MASH) (10-12), suggesting the critical role of methionine metabolism in maintaining liver health. Moreover, MASLD and MASH are associated with increased hepatic expression of the nuclear receptor peroxisome proliferator-activated receptor gamma (*Pparg*) in mice and humans (13-16). Interestingly, we have reported that expression of *Pparg*, in hepatocytes, contributes to the development of diet-induced MASH (17). Specifically, adult-onset, hepatocyte-specific *Pparg* knockout (*Pparg*^ΔHep^) reduced the progression of diet-induced liver injury (hepatocyte ballooning, inflammation, and fibrosis). Some of the protective effects of *Pparg*^ΔHep^ might be due to preservation of methionine metabolism, where expression of hepatic methyltransferases (*Pemt or Bhmt*) is reduced by diet-induced MASH and restored by *Pparg*^ΔHep^ (16,18). Notably, the transcriptional activity of PPAR in the liver is enhanced by fatty acids and also by specific pharmaceutical drugs called thiazolidinediones (TZDs) (19,20). However, it remains to be determined whether the activation of PPARγ in the liver has a significant effect on the regulation of methionine metabolism and, if so, whether it might contribute to the progression of MASLD to MASH.

In this study, we sought to explore the role of hepatocyte *Pparg* in regulating the transcription of genes involved in methionine metabolism in the liver under different metabolic states in which hepatic methionine metabolism is regulated and fatty acid flux to the liver is altered. Notably, fasting increases the expression of hepatic genes involved in methionine metabolism (21-23), and diet-induced MASH may have an impact on the regulation of hepatic methionine metabolism (8). Thus, we have used liver samples from mice that were subjected to 24h fasting, with or without 6h of refeeding. Furthermore, we used liver samples from mice with diet-induced insulin resistance, obesity, and steatosis but without MASH (7). In these metabolic conditions, we have assessed the effect of hepatocyte *Pparg* expression and that of TZD-mediated activation of hepatocyte PPAR on the transcriptional regulation of genes involved in hepatic methionine metabolism.

## Materials and methods

### Mouse studies

Animal studies were conducted with the approval of the University of Illinois Chicago (UIC) and Jesse Brown VA Medical Center (JBVAMC) Institutional Animal Care and Use Committee (IACUC). For these studies, PPARγ floxed mice in a C57B1/6J background were maintained as a homozygous breeding colony (*Pparg*^fl/fl^, Strain B6.129-Pparg^tm2Rev^/J, stock number 004584, Jackson Laboratories, Bar Harbor, ME). *Pparg*^ΔHep^ mice were generated by injection of adeno-associated virus serotype 8 (AAV8) with a thyroxine-binding globulin (TBG)-promoter driving Cre recombinase (AAV8-TBGp-Cre, Penn Vector Core, University of Pennsylvania, and Addgene, Watertown, MA, USA) into the lateral tail vein of *Pparg*^fl/fl^ mice as we previously reported (7,13). An injection of AAV8-TBGp-Null in *Pparg*^fl/fl^ mice was used to generate control mice.

#### Fasting and Refeeding

*Pparg*^fl/fl^ mice were housed in a temperature (22-24°C) and humidity-controlled specific pathogen-free barrier facility (JBVAMC animal facility) with 12h light/ 12h dark (lights on 0600h) and fed a standard chow diet (Formulab Diet 5008, Purina Mills, Richmond, IN) *ad libitum*. At 10-12 weeks of age, *Pparg*^fl/fl^ mice were injected with AAV8-TBGp-Cre or AAV8-TBGp-Null (13). One week after AAV injection, chow-fed control and *Pparg*^ΔHep^mice were fasted for 24 hours before euthanasia by decapitation without anesthesia (food removed at 1200h). A subset of 24h-fasted mice from each group (food removed at 0700h), were then refed with a chow diet at 0700h, and euthanized by decapitation without anesthesia 6h later at 1300h. To determine if fasting and refeeding altered hepatic gene expression from or to a “fed-like” state, liver samples from a previously published study were used in which fed mice were euthanized by decapitation without anesthesia in the post-absorptive state, 4h after food withdrawal at 0700h (13). Blood glucose levels were determined using Blood Glucose Meter and test strips (Accu-Check ®, Roche), and trunk blood was collected in EDTA-coated tubes to obtain plasma and stored at -20°C. Tissues were weighed and rapidly snap-frozen in liquid nitrogen for molecular analyses. All the liver samples used for this study were obtained from littermate mice, which were simultaneously euthanized from 1100h to 1300h.

#### Diet-induced obese mice

To examine the impact of diet-induced insulin resistance, obesity, and steatosis on the expression of hepatic genes involved in the metabolism of methionine in the presence and absence of hepatocyte *Pparg*, liver samples from a previously published study were used (7). In brief, *Pparg*^fl/fl^ mice were housed in a temperature (22-24°C) and humidity-controlled specific pathogen-free barrier facility with 14 h light/ 10 h dark (UIC animal facility). Four-to six-week-old male *Pparg*^fl/fl^ littermate mice were switched from a standard chow diet, to either a low-fat diet (LF, LFD) containing 10% kcal from fat (D12450J, Research Diets, New Brunswick, NJ, USA) or a high-fat diet (HFD) containing 60% kcal from fat (D12492, Research Diets) for 16 weeks. Then, AAV vectors were intravenously injected to generate control and *Pparg*^ΔHep^ mice. One week later, a subset of HFD-fed control and *Pparg*^ΔHep^ mice were switched to a HFD supplemented with 70 mg of rosiglitazone maleate/kg of diet (HFD/TZD, Cat# D18061406, Research Diets) for 6 additional weeks. All the groups were euthanized by decapitation without anesthesia 4h after food withdrawal at 0700h. Livers were weighed and snap-frozen in liquid nitrogen for molecular analyses.

### Mouse primary hepatocytes (MPH)

MPH were obtained from 4 different groups: 1) **CHOW**-chow-fed 13-to 20-week-old male *Pparg*^fl/fl^ mice, 2) **LFD** - male *Pparg*^fl/fl^ mice injected with AAV8-TBGp-Null at 11-14 weeks of age and fed a LFD (D12450J) for 25-26 weeks, 3) **HFC+Fr** - male *Pparg*^fl/fl^ mice that were treated with AAV8-TBGp-Null at 12-13 weeks of age and fed a HFD containing 60% kcal from fat and 2% cholesterol (HFC, D12120101, Research Diets) supplemented with 10% fructose in the drinking water (HFC+Fr) for 24 weeks, 4) **HFC+Fr KO** - male *Pparg*^fl/fl^ mice that were injected with AAV8-TBGp-Cre at 11-13 weeks of age to generate *Pparg*^ΔHep^ and fed a HFC+Fr diet for 25-26 weeks. Mice from these four groups were anesthetized with ketamine/xylazine (100/10 mg/kg) and euthanized by exsanguination during the perfusion of the liver. MPH were isolated as previously described (18) with slight modifications. The liver was perfused using a perfusion system for cell isolation from liver (Harvard Apparatus, Holliston, MA) to control flow rate, and temperature at 37°C. First, we perfused up to 30 ml of Hanks’ Balanced Salt Solution without calcium or magnesium supplemented with EDTA (137 mM NaCl, 5.33 mM KCl, 0.44 mM KH2PO4, 0.33 mM Na2HPO4, 0.5 mM EDTA, pH 7.3) at 3.12 ml/min. Then, 12.5 to 20ml DMEM without sodium pyruvate but supplemented with HEPES and collagenase (4.5 g/l glucose, 110 mM NaCl, 5.33 mM KCl, 44 mM NaHCO3, 0.7 mM Na2HPO4·H2O, 97.67mM MgSO4, 1.8 mM CaCl2, 15 mM HEPES, 25 μg/ml collagenase type I and II Liberase TM, (Sigma Aldrich, St. Louis, MO), pH 7.3) was perfused at 3.12ml/min. When digestion was completed, livers were gently dissociated using cell scrapers in ice-cold DMEM/F-12 supplemented with 10% fetal bovine serum, 2 mM L-glutamine, 1x penicillin-streptomycin. The cell suspension was filtered gently through 70 µm nylon strainers and centrifuged (5min, 100g, 4°C) to pellet the hepatocytes, that were washed three times and centrifuged (5min, 100g, 4°C) with fresh ice-cold complete DMEM/F-12. If cell viability determined by trypan blue exclusion was below 70%, viable hepatocytes were purified by density separation with 45% Percoll (Cytiva, Marlborough, MA) in phosphate-buffered saline buffer by centrifugation (20min, 100g, 4°C, without deceleration). Isolated hepatocytes were resuspended in ice-cold complete DMEM/F-12 and plated at 2*10^5^ cells/well on 12-well plates precoated with type I rat tail collagen (Corning, Corning, NY). After 4 hours at 37°C in a humidified incubator with 5% CO_2_, the medium was replaced with fresh complete DMEM/F12 with and without 1uM rosiglitazone (Sigma-Aldrich), and the cells were incubated for an additional 24 hours at 37°C with 5% CO_2_.

### Hepatic lipid extraction, hepatic and plasma metabolic endpoint assays

Hepatic lipids were extracted using isopropyl alcohol as we previously described (24). Plasma non-esterified fatty acids (NEFA) and plasma and hepatic triglycerides (TG) were measured using colorimetric assays (Fujifilm Wako Diagnostics, Richmond, VA). Plasma insulin levels were determined using a commercial ELISA immunoassay kit (Mercodia, Uppsala, Sweden). All assays were performed according to the manufacturer’s protocols.

### RNA isolation, cDNA synthesis, Real-Time PCR

Hepatic and MPH RNA was extracted using Invitrogen™ TRIzol™ Reagent (Invitrogen, Carlsbad, CA) according to the manufacturer’s protocol. Extracted total RNA was treated with RQ1 RNase-Free DNase (Promega, Madison, WI). DNA-free RNA was then reverse-transcribed into cDNA using the First Strand cDNA Synthesis Kit (Thermo Scientific, Waltham, MA). A Real-Time qPCR reaction was performed using the Brilliant III Ultra-Fast QPCR Master Mix (Agilent Technologies, Santa Clara, CA). Specific primer sequences (Table 1) were used to amplify target genes. Standard curves were generated to determine the number of mRNA copies per sample, as we previously described (24,25). mRNA copy number was adjusted by a normalization factor calculated from the mRNA copy number of at least two separate housekeeping genes (β-actin, and cyclophilin-A mRNA) using GeNorm 3.3 software (24,26).

**Table 1.**
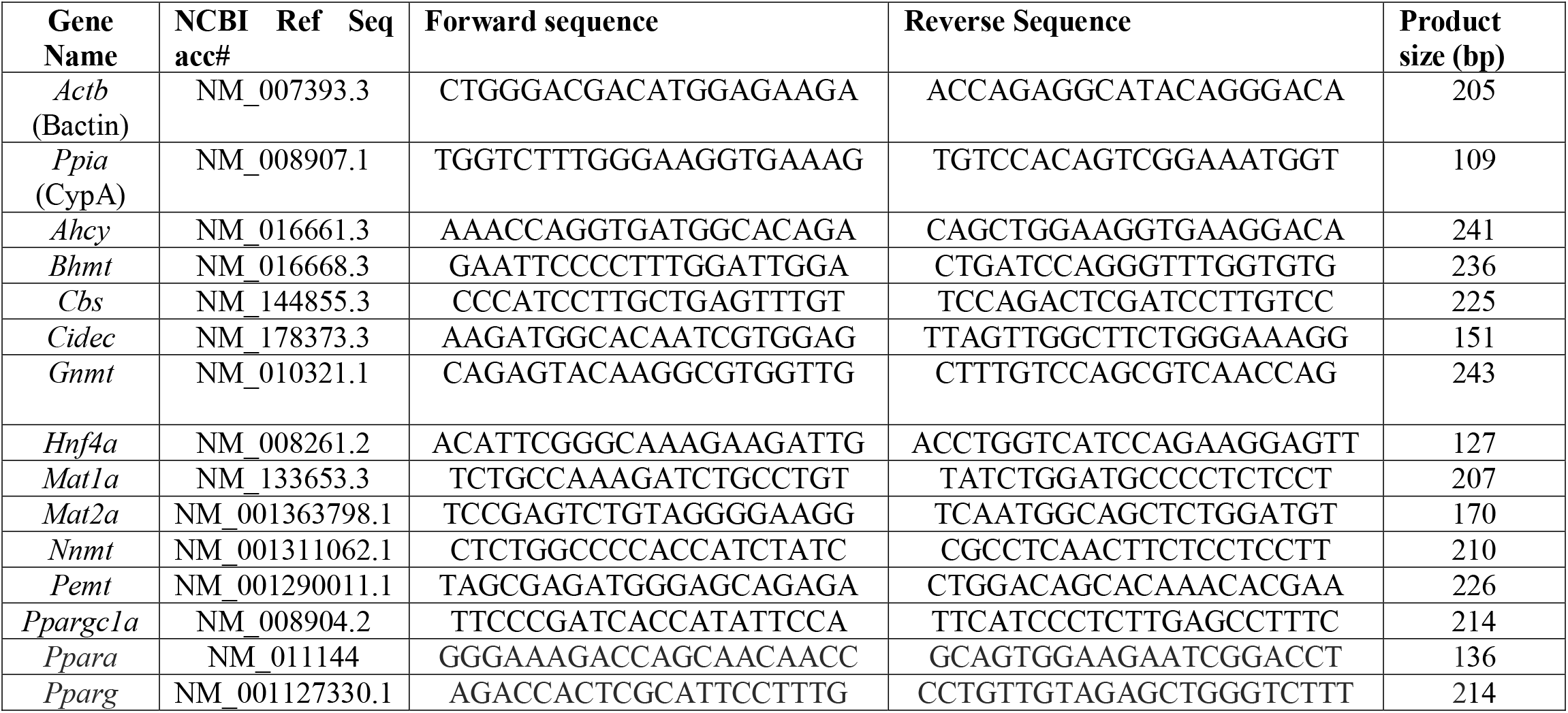
Sequence of qPCR primers used in this study.

### Statistical analysis

Statistical analyses were performed using GraphPad Prism 5 software (GraphPad, La Jolla, CA). Mouse study data were analyzed using a one-way analysis of variance (ANOVA), followed by a Tukey’s post hoc test to compare the effect of fasting and refeeding or the effect of HFD and TZD treatment in the HFD-fed mice. An unpaired Student’s t-test was used to compare the data between control and *Pparg*^ΔHep^ mice under feeding/diet conditions, or the effect of rosiglitazone (TZD) in MPH. Data are presented as mean ± standard error of the mean (SEM), and p-values <0.05 were considered significant for all statistical tests.

## Results

### Effect of fasting and refeeding in the regulation of hepatic methionine metabolism

Since fasting increases hepatic methionine metabolism, we have assessed the effect of *Pparg*^ΔHep^ in a group of fasted and refed mice. Whereas 24h fasting decreased body weight, liver weight, and glucose levels, it increased plasma NEFA levels in control mice (Fig. 1A-F). In this group of control mice, fasting-associated increase in liver TG and decrease in plasma insulin did not reach statistical significance (Fig. 1C, E). Of note, six hours of refeeding with a chow diet did not significantly increase body weight. However, it increased liver weight, glucose, and insulin levels, and decreased the levels of plasma NEFA in control mice as compared to fasted mice (Fig. 1A-F). These effects on liver and plasma endpoints indicate that nutrients were assimilated, and that the mice transitioned from fasting to postprandial conditions. In *Pparg*^ΔHep^ mice, fasting had similar effects to those in control mice in body weight, liver weight, blood glucose, plasma insulin, and NEFA levels (Fig. 1A-F), and it reached statistical significance in liver TG in *Pparg*^ΔHep^ mice (Fig. 1C). Of note, *Pparg*^ΔHep^ increased the levels of plasma insulin and NEFA in refed mice (Fig. 1E-F) without altering body weight or liver weight, which could be associated with some degree of insulin resistance due to the loss of hepatocyte *Pparg* expression.

**Figure 1.**
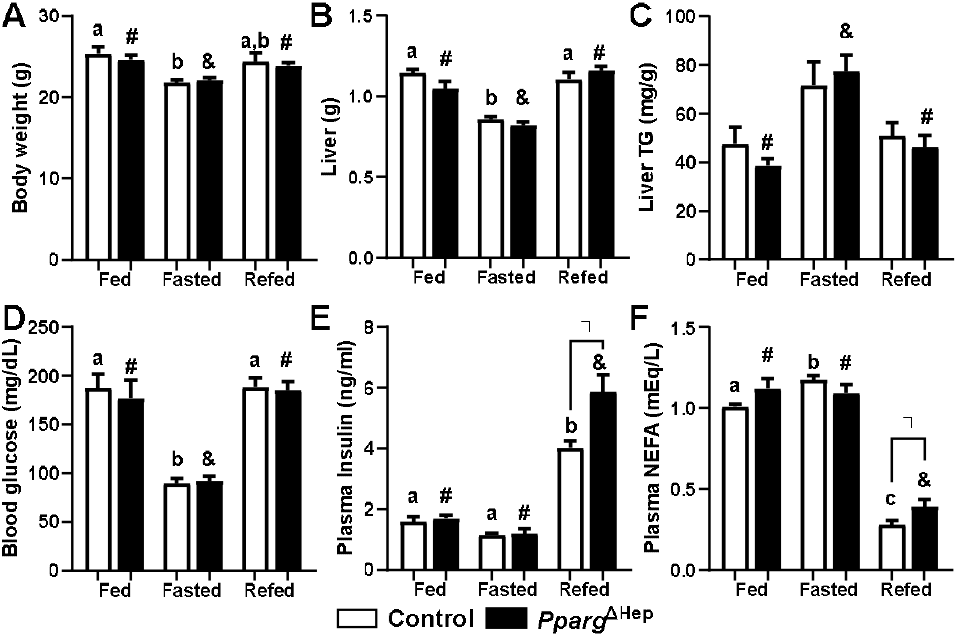
Metabolic effects of fasting and refeeding in control and Pparg^ΔHep^ mice. A) body weight, B) liver mass, C) liver triglicerydes (TG) levels, D) blood glucose levels, E) plasma insulin, and F) non-estrified fatty acids (NEFA) levels of male chow-fed control and *Pparg*^ΔHep^ mice that were euthanized in fed conditions (food removed at 0700h), or after 24h fasting (Fasted, food removed at 1200h), or after 6h refeeding that followed a 24h fasting (Refed, food removed at 0700h the previous day, and added 6h before euthanasia). Data are represented as means ± standard error of the mean. Statistical differences (p<0.05) are indicated by different letters (a, b, c, in control mice) or symbols (#, &, in *Pparg*^ΔHep^ mice). Asterisk indicates statistical differences (p<0.05) between control and *Pparg*^ΔHep^ mice within a feeding state. n=4-6 mice/group. Data of fed mice in Figure 1A-F was previously reported by our group (13).

In this study, 24h fasting significantly increased the hepatic expression of *Mat1a, Mat2a, Gnmt, Nnmt, Ahcy*, and *Bhmt* in control mice (Fig. 2A-G). Only, 6h of refeeding reversed the effect of fasting in the expression of *Mat1a, Mat2a, Nnmt, Ahcy*, and *Bhmt* (Fig. 2A,B,D,F,G), and further reduced the expression of *Gnmt* as compared to that of fed control mice (Fig. 2C). Moreover, refeeding reduced the expression of *Pemt* and *Cbs*, that was not altered by fasting (Fig. 2E, H). Of note, fasting increased the expression of proliferator-activated receptor alpha (*Ppara*), hepatocyte nuclear factor 4 alpha (*Hnf4a*), and peroxisome proliferator-activated receptor gamma cofactor 1 (*Ppargc1a*), which are direct transcriptional activators of methionine metabolism (23,27), whereas it did not alter hepatic *Pparg* expression in control mice (Fig. 2I-L). Most of the hepatic expression of *Pparg* is derived from hepatocytes, and thereby the expression of hepatic *Pparg* is dramatically reduced in *Pparg*^ΔHep^ mice as compared to control mice (Fig. 2I) as we reported previously (13). However, fasting increased the expression of hepatic *Pparg* in *Pparg*^ΔHep^ mice, suggesting there is an effect of fasting on the expression of *Pparg* in non-hepatocyte cells. Only six hours of refeeding decreased the expression of *Ppara, Hnf4a*, and *Ppargc1a* (Fig. 2J-L), and that downregulation was associated with the reduced expression of genes involved in methionine metabolism in the liver (Fig. 2A-H). Overall, the loss of hepatocyte *Pparg* expression did not alter the regulation of the hepatic genes involved in methionine metabolism in fasted and/or refed mice, which indicates that in these conditions, the expression of hepatocyte *Pparg* is not essential to regulate the methionine cycle. However, *Pparg*^ΔHep^ reduced the expression of *Ppara* in refed mice, and that of *Hnf4a* in fasted and refed mice.

**Figure 2.**
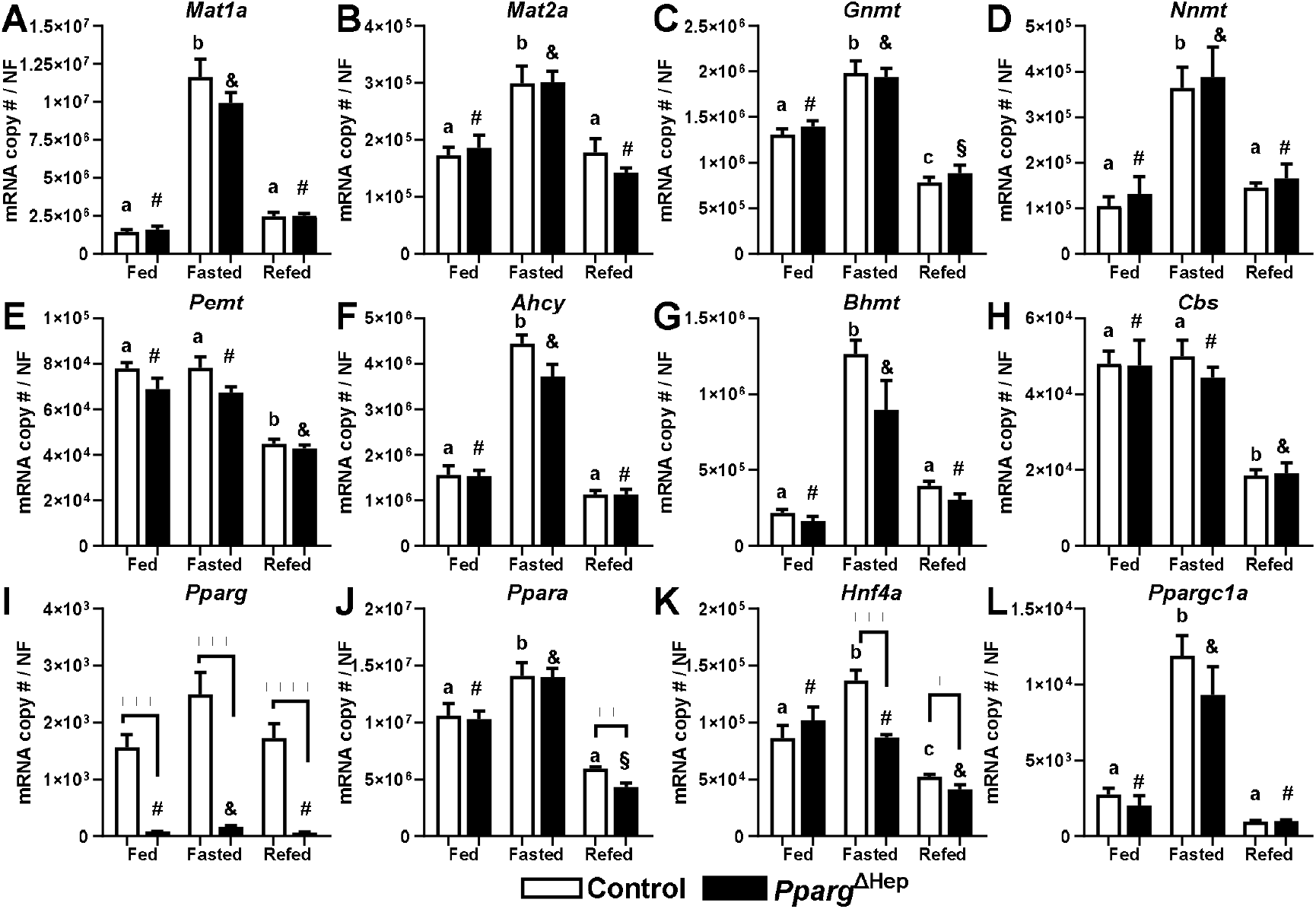
Effects of fasting and refeeding on the hepatic expression of control and Pparg^ΔHep^ mice. Expression of A) *Mat1a*, B) *Mat2a*, C) *Gnmt*, D) *Nnmt*, E) *Pemt*, F) *Ahcy*, G) *Bhmt*, H) *Cbs*, I) *Pparg*, J) *Ppara*, K) *Hnf4a*, and L) *Ppargc1a* in male chow-fed control and *Pparg*^ΔHep^ mice that were euthanized in fed conditions (food removed at 0700h), or after 24h fasting (Fasted, food removed at 1200h), or after 6h refeeding that followed a 24h fasting (Refed, food removed at 0700h the previous day, and added 6h before euthanasia). Data are represented as the average of mRNA copy number per sample normalized with a normalization factor (NF) ± standard error of the mean. Statistical differences (p<0.05) are indicated by different letters (a, b, c, in control mice) or symbols (#, &, §, in *Pparg*^ΔHep^ mice). Asterisks indicate statistical differences (p<0.05) between control and *Pparg*^ΔHep^ mice within a feeding state. n=4-6 mice/group. **, p<0.01. ***, p<0.001. ****, p<0.0001. Data of fed mice in Figure 2I was previously reported by our group (13).

### Effect of diet-induced obesity and PPARγ activation in the regulation of hepatic methionine metabolism

We previously reported that hepatocyte *Pparg* expression is negatively associated with the regulation of methionine metabolism in conditions of advanced MASLD/MASH (18). Here, we have assessed the PPARγ-dependent effects of diet-induced obesity and TZD-mediated activation of PPARγ in control and *Pparg*^ΔHep^ mice, that were previously reported by our group (7). In our previous studies, we showed that HFD increased body weight, liver weight, liver TG, and plasma insulin levels in control mice, and TZD increased body weight and liver steatosis while lowering insulin levels due to its insulin-sensitizing effects (7). Regarding the effects of diet-induced obesity on the control of methionine metabolism, HFD only reduced the expression of hepatic *Pemt* (Fig. 3A-H) in control mice. In diet-induced obese control mice, TZD reduced the expression of hepatic *Bhmt* and *Cbs* (Fig. 3G,H). We also have reported that *Pparg*^ΔHep^ decreased liver weight and steatosis in HFD-fed mice (7). In *Pparg*^ΔHep^ mice, HFD did not alter the expression of hepatic genes involved in methionine metabolism, but TZD increased the expression of hepatic *Cbs* (Fig. 3H). Interestingly, the comparison between TZD-treated control and *Pparg*^ΔHep^ mice shows that expression of *Mat1a, Gnmt, Nnmt, Pemt, Ahcy, Bhmt*, and *Cbs* was increased in *Pparg*^ΔHep^ mice treated with TZD (Fig. 3A-H). Furthermore, the expression of *Pemt* was increased in HFD-fed *Pparg*^ΔHep^ mice as compared to HFD-fed controls (Fig. 3E). These data might indicate that expression of *Pparg*, or TZD-mediated activation of PPARγ in hepatocytes, could negatively regulate the expression of genes involved in methionine metabolism in the liver. As we previously reported (7), the expression of hepatic *Pparg* was significantly increased by HFD in control mice (Fig. 3I). Interestingly, neither HFD nor the TZD treatment in HFD significantly reduced the expression of the regulators of methionine metabolism: hepatic *Ppara, Hnf4a*, or *Ppargc1a* in control mice (Fig. 3J-L). However, HFD reduced the expression of *Ppargc1a* in *Pparg*^ΔHep^ mice, as compared to that of LFD-fed *Pparg*^Δhep^ mice (Fig. 3L). Moreover, TZD increased the expression of hepatic *Hnf4a* in HFD-fed *Pparg*^Δhep^ mice (Fig. 3K). In addition, when we compared TZD-treated control and *Pparg*^ΔHep^ mice, the expression of *Hnfa* and *Ppargc1a* was increased in *Pparg*^ΔHep^ mice treated with TZD (Fig. 3K-L), and that could be associated with a positive regulation of most of the genes involved in hepatic methionine metabolism (Fig. 3A-H).

**Figure 3.**
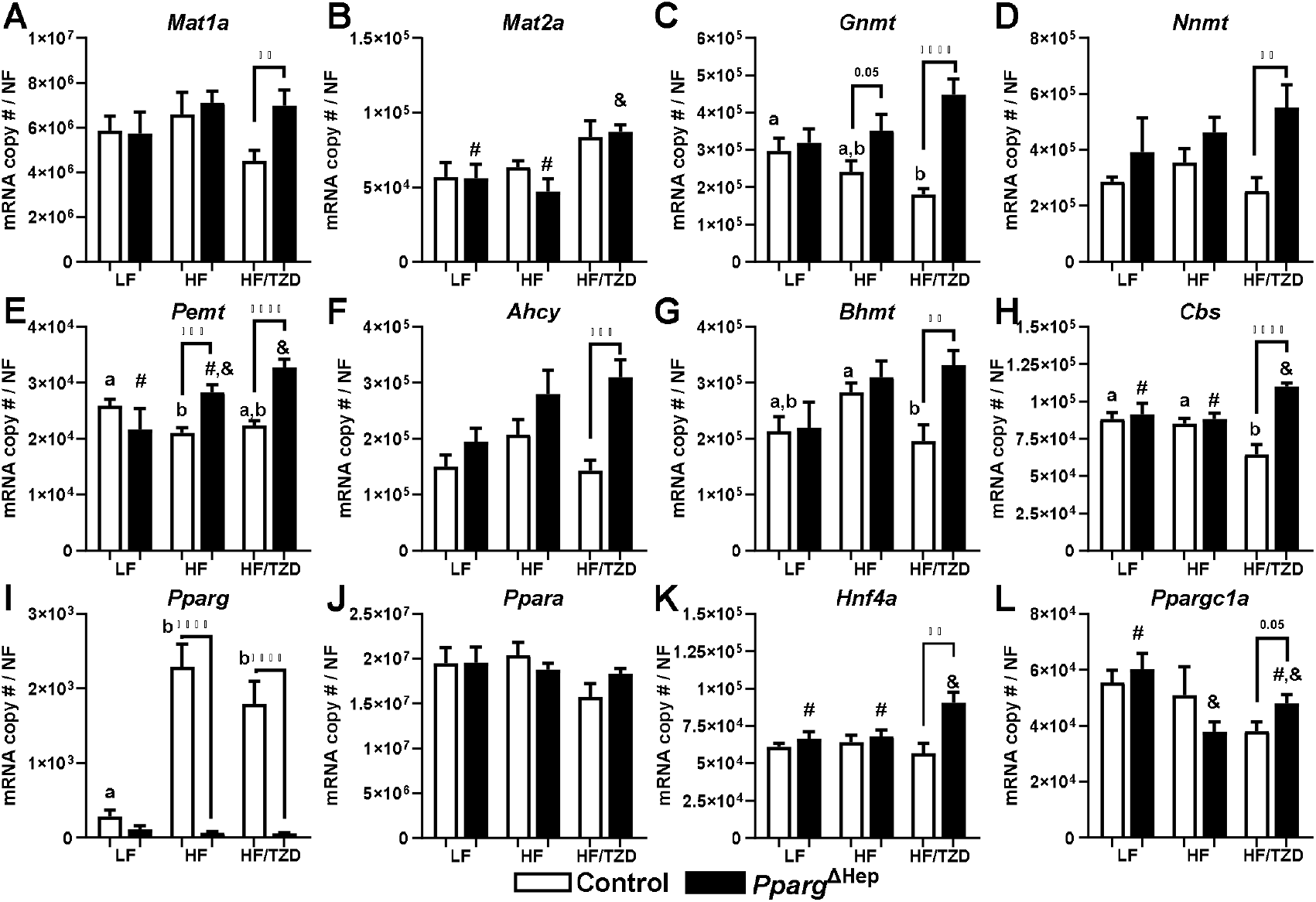
Effects of high-fat diet-induced obesity, and TZD-mediated activation of PPARγ in obese mice on the hepatic expression of control and Pparg^ΔHep^ mice. Expression of A) *Mat1a*, B) *Mat2a*, C) *Gnmt*, D) *Nnmt*, E) *Pemt*, F) *Ahcy*, G) *Bhmt*, H) *Cbs*, I) *Pparg*, J) *Ppara*, K) *Hnf4a*, and L) *Ppargc1a* in male control and *Pparg*^ΔHep^ mice that were fed a LF or HF diet for 23 weeks. A subset of HF-fed control and *Pparg*^ΔHep^ mice were treated with TZD for the last 6 weeks of diet (HF/TZD). Data are represented as the average of mRNA copy number per sample normalized with a normalization factor (NF) ± standard error of the mean. Statistical differences (p<0.05) are indicated by different letters (a, b in control mice) or symbols (#, &, in *Pparg*^ΔHep^ mice). Asterisks indicate statistical differences (p<0.05) between control and *Pparg*^ΔHep^ mice within a feeding state. n=7-9 mice/group. **, p<0.01. ***, p<0.001. ****, p<0.0001.

Overall, the positive effects of TZD in the expression of *Mat1a, Gnmt, Nnmt, Pemt, Ahcy, Bhmt*, and *Cbs* and their regulators *Hnf4a* and *Ppargc1a* may be indicative of a negative association between hepatocyte *Pparg* and the regulation of hepatic methionine metabolism. As we previously reported (7), TZD-treated diet-induced obese *Pparg*^ΔHep^ mice show a significant improvement in liver health that could be associated with enhanced effects of TZD-mediated insulin sensitization in the regulation of hepatic methionine metabolism.

To further investigate the role of hepatocyte PPARγ in the regulation of methionine metabolism under diet-induced metabolic stress, we isolated MPH from metabolically healthy 4.5-month-old chow-fed and 9-month-old LFD-fed mice, and from obese, metabolically unhealthy 9-month-old HFC+Fr-fed control and *Pparg*^ΔHep^ mice. In these MPH, we assessed the effect of TZD-mediated activation of hepatocyte PPARγ in the regulation of some hepatocyte genes. Notably, TZD increased the expression of the PPARγ-target gene cell death-inducing DFFA-like effector c (*Cidec*) in MPH of chow-fed mice, and HFC+Fr-fed control mice (Fig. 4A), similar to what we published previously with a higher dose of TZD (18). Interestingly, TZD reduced the expression of *Bhmt* in MPH from chow-fed mice, and HFC+Fr-fed control mice (Fig. 4B). Furthermore, TZD reduced the expression of *Cbs* in chow-fed mice, and trend to reduce that in HFC+Fr-fed control mice (Fig.4C). The TZD-mediated regulation of *Cidec, Bhmt*, and *Cbs* was not observed in MPH obtained from HFC+Fr-fed *Pparg*^ΔHep^ mice. These data suggest that TZD-mediated activation of PPARγ in hepatocytes may directly reduce the expression of *Bhmt* and *Cbs*, and that could alter hepatocyte methionine metabolism.

**Figure 4.**
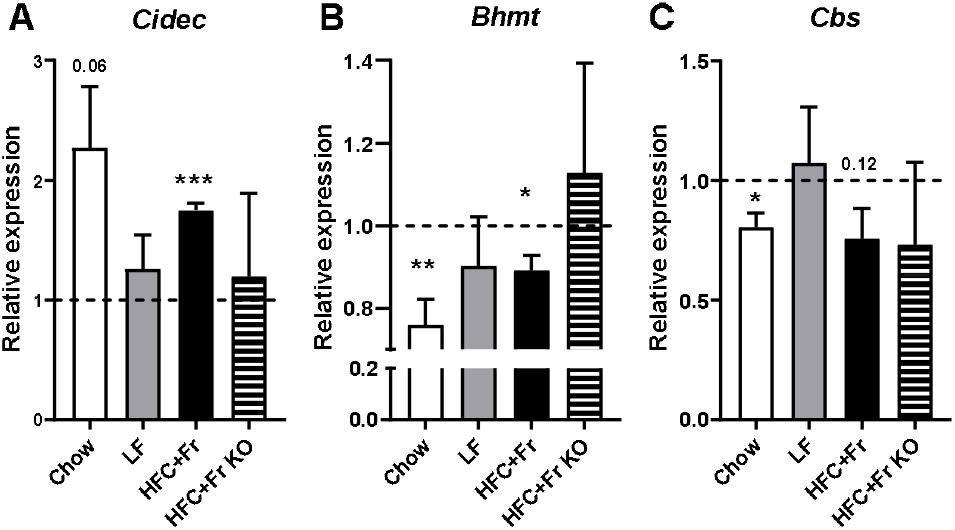
Effects of TZD-mediated activation of hepatocyte PPARγ on the gene expression of mouse primary hepatocytes. Expression of A) *Cidec*, B) *Bhmt*, and C) *Cbs* in mouse primary hepatocytes of chow-fed Pparg^fl/fl^ mice, LF-fed control, HFC+Fr-fed control, and HFC+Fr-fed *Pparg*^ΔHep^ (KO) mice. Data are represented as the average of relative gene expression values ± standard error of the mean against their vehicle (Veh)-treated mouse primary hepatocytes (set as a discontinued line at 1). Asterisks indicate statistical differences between TZD-treated MPH vs vehicle-treated MPH. *, p<0.05. **, p<0.01. ***, p<0.001. n=3-4/group.

## Discussion

Most SAM-dependent methylation reactions take place in the liver and play a key role in maintaining liver health (1,6,28,29). The enzymes that generate SAM, as well as many SAM-dependent methyltransferases, are highly expressed in hepatocytes (28-30). In hepatocytes, cellular stress and fatty acids activate key transcription factors and nuclear receptors to regulate the expression of the genes involved in the methionine cycle and to maintain cell health. Fasting, refeeding, and diet-induced steatosis can induce cellular stress in hepatocytes and alter the flux of fatty acids to the liver which activates key transcription factors involved in the regulation of methionine metabolism. In this study, we have reproduced and extended the knowledge of the expression of genes involved in methionine metabolism in the liver under fasting, refeeding, and diet-induced steatosis, and show that hepatocyte *Pparg* is not required for the regulation of methionine metabolism genes in the context of fasting and refeeding, but it plays a negative role in the regulation of key methyltransferases in conditions of obesity and insulin resistance.

It is known that fasting enhances hepatic methionine metabolism by increasing the expression of *Mat1a* and major methyltransferases (21-23). Our study shows that both *Mat1a* and *Mat2a* expression is increased in fasted livers, although previous reports reported that *Mat2a* was not increased by fasting (22,23). Of note, Capelo-Diz *et al*. reported that increased methylation reactions during fasting may be required to prevent liver damage due to increased metabolism in fasting conditions and to maintain mitochondrial oxidative capacity and ATP production (21). Here, we report that fasting increases the expression of major SAM-dependent methytransferases, including *Gnmt* and *Nnmt*, which may consume the SAM produced by MAT1A activity in the hepatocytes during fasting. Also, we observed that fasting increases the expression of hepatic *Ahcy* and *Bhmt*, which will convert SAH to homocysteine and back to methionine (23,31). Increased activity of SAM-dependent methyltransferases may lead to an increase in homocysteine, which can induce endoplasmic reticulum stress and dysregulate lipid metabolism (32), so the upregulation of *Bhmt* will also maintain liver health. Interestingly, our study shows that fasting does not increase the expression of hepatic *Pemt* and *Cbs*, which might indicate that use of SAM for hepatic VLDL or use of homocysteine for glutathione production during fasting does not require increased PEMT or CBS activity, or that these enzymes may not significantly deplete SAM and homocysteine pools during fasting, respectively. Overall, we have reproduced the effect of fasting in the control of *Mat1a, Ahcy, Bhmt*, and *Cbs* as previously reported by others (23,31), and extended it to additional genes (*Mat2a, Gnmt, Nnmt*, and *Pemt*). In addition, our study shows that refeeding (postprandial phase) reverses the fasting effect on most of these genes and reduces the basal expression of hepatic *Pemt* and *Cbs*, which might impact VLDL secretion rate and glutathione production. The regulation of the genes involved in methionine metabolism may be produced by activation of *Ppargc1a* and *Hnf4a* during fasting, which are known to activate the expression of some of these genes such as *Mat1a* and *Bhmt* (23,27). Our study also reveals that in lean mice, despite the fasting-associated influx of fatty acids that might activate hepatocyte PPARγ (33), the expression of *Pparg* in hepatocytes does not affect the fasting/refeeding regulation of genes involved in methionine metabolism.

Diet-induced obesity also regulates hepatic methionine metabolism (8). High-fat diets increase body weight, plasma insulin levels, and liver steatosis in C57BL/6J male mice (7), and these diets may promote the onset of MASLD. MASLD may progress to MASH, which is strongly associated with the negative regulation of hepatic methionine metabolism by affecting the production of SAM, expression of SAM-dependent methyltransferases, and the remethylation or transulfuration of homocysteine (8,16,18). Specifically, a western diet with 42% Kcal from fat reduces the expression of hepatic *Mat1a, Mat2a, Gnmt*, and *Cbs* gene in female C57Bl/6J mice (8), a high fat diet with 45% Kcal from fat increases the expression of hepatic *Bhmt2* and decreases that of *Pemt* and *Cbs* in male C57Bl/6J mice (34), and a high fat diet with 60% Kcal from fat increases hepatic *Bhmt* and trended to reduce *Cbs* gene expression in male C57Bl/6N mice (35). We extended these observations in male C57Bl/6J mice fed a high-fat diet with 60% Kcal from fat (7) in which only hepatic *Pemt* expression was reduced, which could negatively impact VLDL secretion rate and promote steatosis (11). Furthermore, we observed that TZD-mediated insulin-sensitizing effects and activation of hepatocyte PPARγ reduced the expression of hepatic *Bhmt* and *Cbs*, which could negatively impact the remethylation and transulfuration of homocysteine. The negative regulation of *Bhmt* and *Cbs* was *Pparg*-dependent and could contribute to the increase of homocysteine in hepatocytes (8,16,18) and that might induce cellular stress in the liver (32). Noteworthy, this *Pparg*-dependent *in vivo* effect was not associated with a downregulation of *Ppargc1a, Hnf4a*, or *Ppara* expression that are known to increase *Bhmt* expression. In fact, this effect seems to be direct as mouse primary hepatocytes treated with rosiglitazone showed a decreased expression of *Bhmt* and *Cbs*. Moreover, the TZD-mediated insulin-sensitizing effects (7) had a positive effect in *Pparg*^ΔHep^ mice because the expression of *Hnf4a*, and *Cbs* is upregulated above that of HF-fed *Pparg*^ΔHep^ mice. Taken together, this data show that pharmacological activation of hepatocyte PPARγ negatively impacts methionine metabolism in hepatocytes, which could reduce the ability of the cell to mediate transmethylation reactions that use SAM, and to reduce the levels of homocysteine that is processed by BHMT and CBS to prevent liver damage.

In conclusion, our study describes the tight regulation of hepatic methionine metabolism between fasting and feeding cycles that controls the expression of genes involved in the synthesis of SAM, the transmethylation of SAM, and the final remethylation or transsulfuration of homocysteine, where these changes are independent of hepatocyte *Pparg*. Although diet-induced obesity does not impose a strong regulation of hepatic methionine metabolism, the expression of *Pemt* was reduced in the livers of obese mice, in a PPARγ-dependent manner. Furthermore, the activation of hepatocyte PPARγ with an exogenous agonist downregulated the expression of key genes involved in the remethylation or transsulfuration of homocysteine. Since the progression of MASLD is associated with increased homocysteine levels (8,16,18,36-38), and homocysteine increases the risk of advanced hepatic fibrosis in patients with alcoholic liver disease (39,40), hepatocyte PPARγ activation may have negative consequences for liver health, and its activity in hepatocytes should be considered when PPAR agonist are used in the clinic.

